# PACAS: pairwise comparisons of aligned subsequences

**DOI:** 10.1101/2021.02.01.428731

**Authors:** Kenneth DeMonn, Christopher L. E. Powell, Fabia U. Battistuzzi

## Abstract

Evaluation of conservation levels of alignments is a pivotal step for evolutionary analyses. Commonly used software to analyze alignments produce either site-by-site or overall measures of conservation without the ability of performing comparative analyses among sub-regions of the alignment. We have developed a stand-alone alignment analyzer (<u>Pa</u>irwise <u>C</u>omparisons of <u>A</u>ligned <u>S</u>ubsequences) that uses user-defined sub-regions to report various measures of conservation among pairs of sequences and between the sub-regions and the rest of the alignment. We then used this method to analyze the repetitive regions (low complexity regions; LCRs) of two genes in Plasmodium: AMA1 and CSP. We found that LCRs are not always associated with high entropy (low conservation) suggesting that some LCRs may be less likely to mutate than others.

**Availability and Implementation:** PACAS is publicly available at
https://github.com/BlabOaklandU/PACAS

**Contact:** Fabia U. Battistuzzi

**Supplementary information:** n/a

## Introduction

Genomic sequences are represented by patterns of nucleotide or amino acid strings with different levels of conservation across genes and species. While evolutionary units of sequences are at the level of single residues (e.g., nucleotides), functional units are often represented by high-order patterns composed of multiple residues in a variety of arrangements (e.g., periodic motifs). These can evolve differently depending on the species and the gene being analyzed. There are many algorithms that analyze genomic sequences and they can be broadly classified into two categories based on the type of comparisons they perform: (i) multi-sequence and (ii) within-sequence comparisons. Belonging to the first category are software such as the various implementations of Clustal (Sievers *et al*., 2011; Larkin *et al*., 2007; Chenna *et al*., 2003), which produces multi-sequence alignments. These alignments can then be analyzed to identify conserved *vs*. variable regions of genes providing many valuable information including domain boundaries, functionally important sites, and evolutionary rates. The second category, instead, focuses on pattern recognition and includes software like Tandem Repeat Finder and SEG (Benson, 1999; Wootton and Federhen, 1993). These identified sub-regions of a sequence are compositionally distinct from other parts allowing for the evaluation of gene structures, origin of new functions, and variation in genome complexity (e.g., low complexity regions). Tools within each of these categories are fundamental for evolutionary analyses as they provide a comparative framework to observe modifications through time. Unfortunately, bridging the results of these two sets of tools to analyze simultaneously among-species and within-gene evolutionary patterns, can be difficult. For example, alignments can be easily queried for information regarding conserved *vs*. non-conserved sites, but obtaining basic information about high-order structures, such as repetitive sequences, usually requires additional steps.

In addition to the need for a unifying tool for multi-sequence and within-sequence comparisons, there is also a need to develop a graphical viewer that will allow easy interpretation of the data. The most similar tool to the one we developed is a recent software (Mutual Information Analyzer) that provides a user-friendly platform to analyze multiple sequence alignments (Lichtenstein *et al*., 2015). However, it lacks the option of analyzing user-defined sub-regions, which is the focus of PACAS.

Thus, to address this need, we created a new application for the comparison of user-defined sub-regions within an alignment. These comparisons facilitate the identification of changes in complexity and length of the sub-regions across sequences and allow statistical testing of the significance of observed changes. The software, called PACAS (<u>Pa</u>irwise <u>C</u>omparisons of <u>A</u>ligned <u>S</u>ubsequences), is distributed as lightweight Perl and Python scripts that can be used with user-created input files (alignment and sub-regions parameter file). It can also work directly with outputs from the program SEG, a commonly used software to identify low complexity regions (LCRs). It runs in multiple environments (Windows, Mac, and Unix) and can be used manually or integrated for batch processing. PACAS produces csv tables and also graphic representations of the comparisons among sequences (Fig. 1). We applied this new method to the analysis of LCRs in two genes important for pathogenicity in Plasmodium: AMA1 (Apical Membrane Antigen 1), which is involved in erythrocyte invasion, and CSP (Circumsporozoite protein), which is required for hepatocytes invasion. Because both of these genes are vaccine candidates understanding their evolutionary patterns is key to efforts to eradicate malaria.

**Figure 1.**
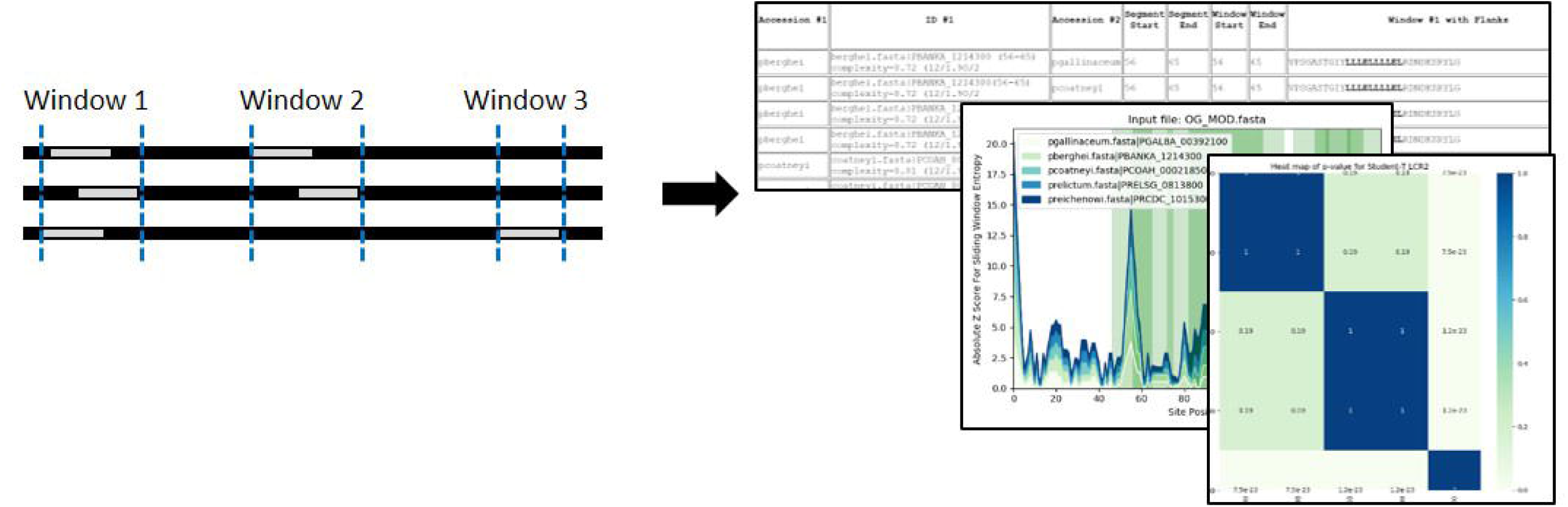
Schematic representation of PACAS. Starting from user-defined sub-regions of an alignment, merged windows are obtained. These windows are then compared in pairs of sequences and against the rest of the alignment. Results are produced in tabular and graphic formats and include a statistical evaluation of the results.

## Methods

The goal of PACAS is to perform comparisons of sub-regions of aligned sequences provided by the user. These sub-regions are compared in pairs to each other and also to the rest of the alignment. These comparisons produce basic sequence information (matches, mismatches, and gaps) and site-by-site sequence conservation analyses using the JSD metric (Capra and Singh, 2007). This information can be used to determine the evolutionary paths of sub-regions relative to the rest of the sequences, a task that can lead to discoveries about domain evolution, among-site rate variation, and other hidden substructures within a gene. Initial user input requires two types of information: (1) a list of sub-regions to compare and (2) an alignment, each specified in its own file. The files are plain text or based on the SEG format for sub-regions and the fasta format for the alignment. Sub-regions files require sequence identifiers and positions within the alignment while alignment files use the same sequence identifiers and a sequential (non-interleaved) format.

The *first* step in PACAS is to identify all sub-region boundaries within the alignment for each sequence. In the *second* step, changes in location of the same sub-region in different sequences is taken into account to create merged “windows”. In this step, if a sub-region in one sequence is found to overlap with another one in a different sequence, its boundaries are expanded in one or both directions to include both sub-regions. The process is repeated iteratively until all sub-regions are evaluated. In the *third* step, after all segments have been assigned to a merged window, the user-input values for flanking regions are applied. *Finally*, once the final windows and flanking regions have been identified, the conservation measures are calculated. For each window and flanking region, aligned sequences are compared in pairs to produce the number of matches, mismatches, gaps, and gap-matches that are reported in a tabular format. The whole sequence is also analyzed for overall conservation and the identified windows are highlighted for easy comparisons. These conservation values are also used for each analyzed window to perform a t-test comparison between pairs. The report of tabular, visual, and statistical results facilitates easy comparisons between sub-regions and their background sequence, which can be used to identify unique evolutionary patterns in different regions of a gene.

From PlasmoDB we obtained ClustalOmega alignments for AMA1 and CSP. For CSP we used all sequences available (*P. relictum, P. falciparum 3D7, P. gaboni, P. vivax, P. yoelii, and P. ovale*) while for AMA1 we selected sequences from within the Laverania group (*P. adleri, P. blacklocki, P. billcolinsi, P. reichenowi, P. facilarum 3D7 and IT, and P. praefalciparum*) using *P. vivax* as an outgroup for comparison (Table 1) (Aurrecoechea *et al*., 2009). These alignments were then analyzed with SEG to identify LCRs (windows size 12, complexity thresholds 1.9 and 2.2) (Wootton and Federhen, 1993; Wootton, 1994). The outputs were used as the starting point for PACAS.

**Table 1.**
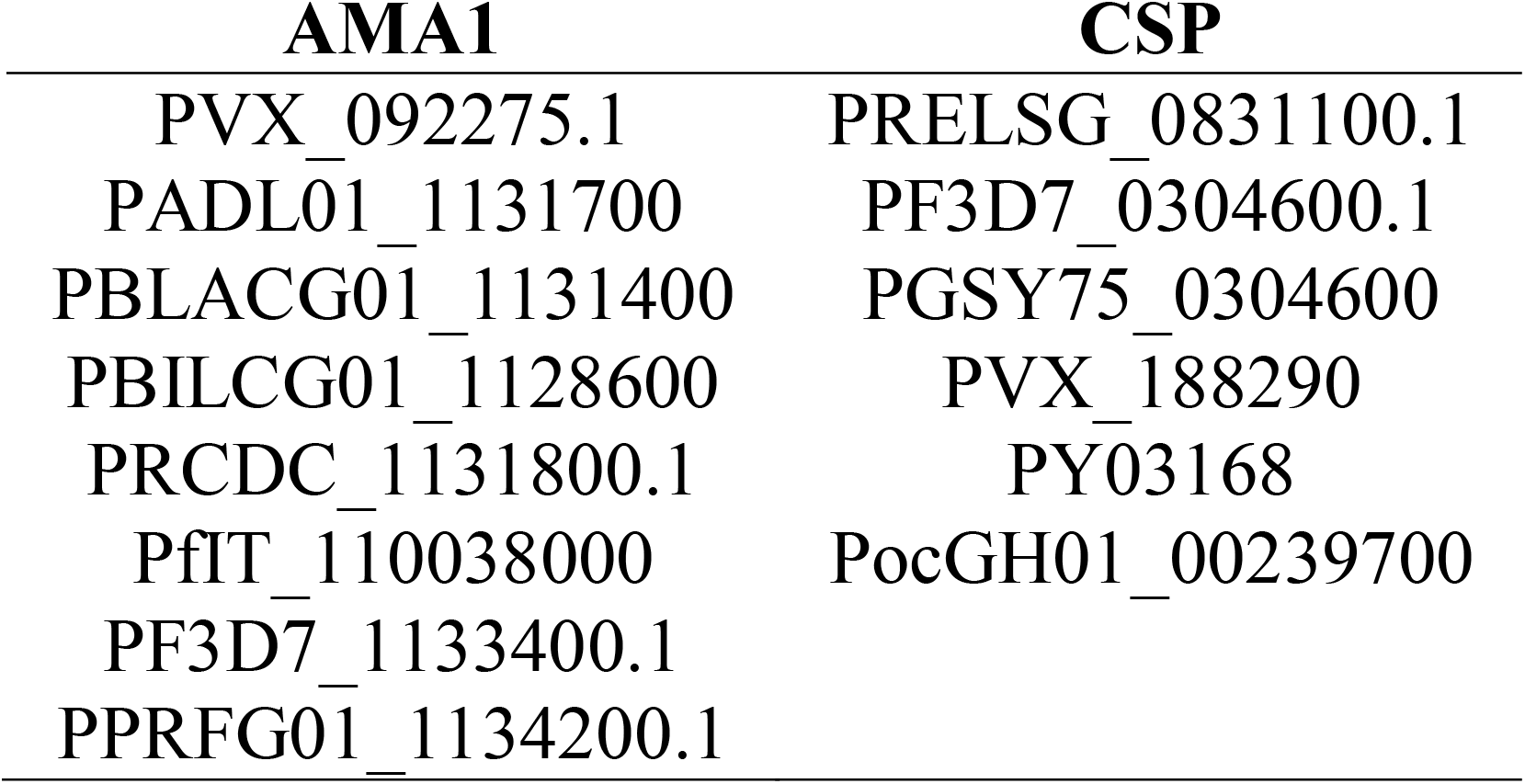
Accession numbers used in the analyses of AMA1 and CSP.

| **AMA1** | **CSP** |
| --- | --- |
| PVX_092275.1 | PRELSG_0831100.1 |
| PADL01_1131700 | PF3D7_0304600.1 |
| PBLACG01_1131400 | PGSY75_0304600 |
| PBILCG01_1128600 | PVX_188290 |
| PRCDC_1131800.1 | PY03168 |
| PfIT_110038000 | PocGH01_00239700 |
| PF3D7_1133400.1 |  |
| PPRFG01_1134200.1 |  |

## Results and Discussion

PACAS is a lightweight software that produces many pairwise comparisons in just a few seconds. We used it to analyze the LCR regions of two genes, AMA1 and CSP, that are important for Plasmodium pathogenicity, being involved in the red blood and liver cycles of the infection, respectively (Healer *et al*., 2005; Ménard *et al*., 1997). Both genes were found to have multiple LCRs that, once merged, resulted in two windows each. Despite previous evidence that LCRs are fast evolving regions (Haerty and Golding, 2011), we found that there is variability in the polymorphism of these regions between and within genes. In AMA1, the first region is clearly associated with a peak in polymorphisms while the second region, albeit being the second highest peak, is less than half as variable as the first one (Fig. 2A). Similarly, the first region in CSP has lower levels of polymorphism, compared to the second region and compared to both regions in AMA1 (Fig 2B, note the difference in scale of the y-axis). Moreover, the second region of CSP, which is the one targeted by the malaria vaccine RTS,S, shows 2-fold variation in entropy, suggesting that sites within the same region are exposed to different evolutionary pressures (Cohen *et al*., 2010).

**Figure 2.**
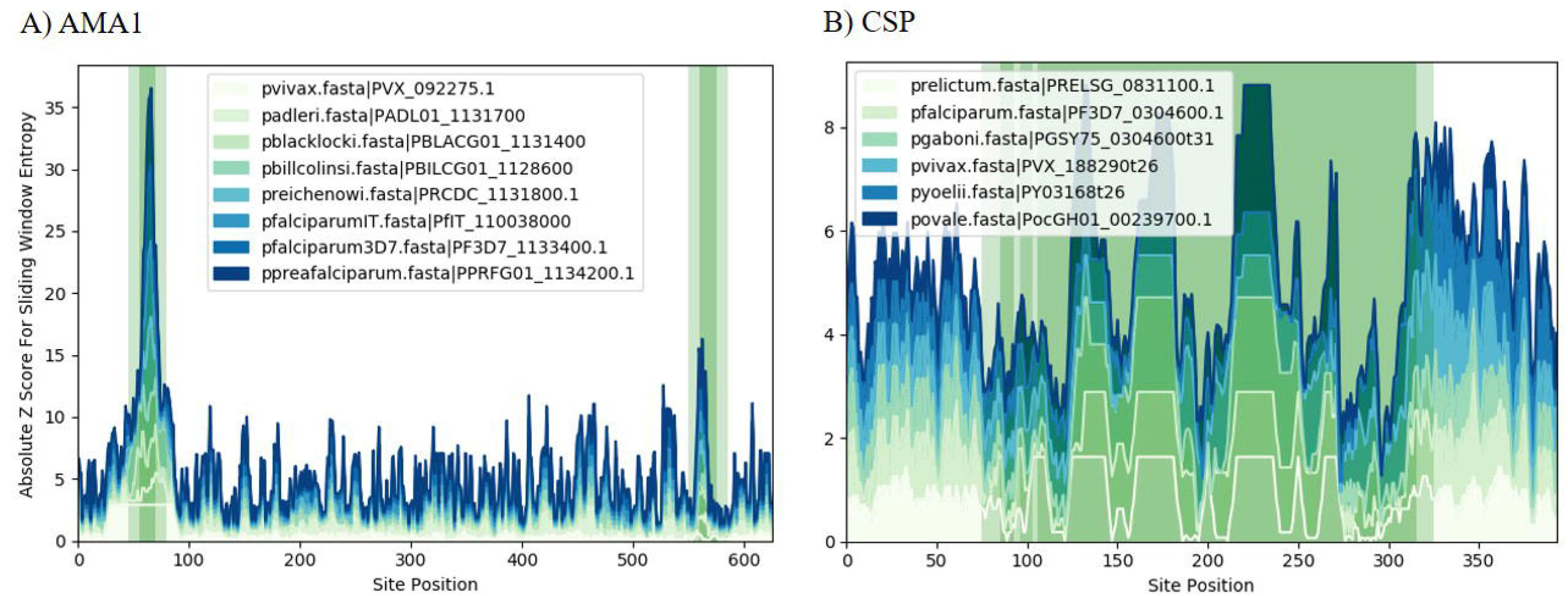
Entropy plots for AMAl (panel A) and CSP (panel B) for selected species in Plasmodium. Peaks in the graph correspond to highly polymorphic regions. Green and shaded green bars are the merged windows and the user-specified flanking regions, respectively.

While the reasons for these differences are still unknown, this analysis shows that direct comparisons of conservation of LCRs within a gene and among species is an effective and easy way to obtain information about possible regions of interest. For example, LCR2 in CSP overlaps with a known epitope suggesting that it may be possible to expand the target region to a larger section with similar properties. At a finer evolutionary level, the methodology used in PACAS was also used to analyze multiple LCRs in multiple strains of *P. vivax* and its closest relatives (Chaudhry *et al*., 2018), which led to the ancestral state reconstruction of these regions over time. All these analyses show how the ability to easily perform comparative genomics of sub-regions within an alignment is a useful tool to reconstruct evolutionary histories of repetitive regions, domains, and other unknown sub-structures of a gene.

## Acknowledgments

We thank Douglas Phelan, Sophia Chaudhry, and Sara Dadashzadeh for preliminary programming and testing of this software.

## Funding

This work was supported by funds from Oakland University and from the National Institute of Health (R15GM121981) to FUB.

## Notes

https://github.com/BlabOaklandU/PACAS

### Competing Interest Statement

The authors have declared no competing interest.

